# InDels in an intronic region of gene *Ccsmd04* coding for dormancy/auxin-associated protein controls sterility mosaic disease resistance in pigeonpea

**DOI:** 10.1101/2025.06.19.660552

**Authors:** Sagar Krushnaji Rangari, Namita Dube, Vinay Sharma, Sunil S Gangurde, Mamta Sharma, Prakash I Gangashetty, Rachit K Saxena, Abhinav Moghiya, Vinay K Sharma, Karma L Bhutia, Ravi Kant, Mahendar Thudi, Satheesh Naik SJ, Aditya Pratap, Girish P Dixit, Sean Mayes, Manish K Pandey

## Abstract

Sterility mosaic disease presents a significant challenge to pigeonpea cultivation in the Indian subcontinent, potentially leading to total crop failure. The development of diagnostic molecular markers for SMD resistance can aid in improving SMD-resistant varieties. In this context, a QTL-seq approach identified genomic regions associated with SMD resistance using a recombinant inbred line generated from ICP8863 × ICPL87119. In total, 6,105 high-confidence variants were identified in the genomic region, namely, *smdCc04* based on the delta SNP index. A genomic region *smdCc04* on chromosome Cc04 spans 3.2 Mb (9.3 - 12.5 Mb) comprised of 6 missense variants and eight indels. A total of 211 candidate genes were identified from this region. 1 bp insertion, 21 bp insertion, 9 bp deletion, and 3 bp insertion at different intronic positions in 22 susceptible line leads to downregulation of *Dormancy/auxin associated protein* (*Ccsmd04*) resulting in loss of signaling in disease resistance pathways. The identified sites recognize four important disease and plant growth related transcription factors. A total of 4 Indels and 8 SNPs were validated from *smdCc04* genomic regions using whole genome re-sequencing data and KASP genotyping on resistant and susceptible pigeonpea lines respectively. These markers will be used in pigeonpea breeding programs.

## Introduction

Pigeonpea or red gram (*Cajanus cajan* L.) emerging as a climate-resilient legume due to its ability to withstand a wide temperature range from 8.2°C to 43°C and its ability to grow with a limited annual rainfall of 65 cm [1–3]. It serves as an important source of protein and considered a ‘life savior crop’ in Asia and Africa’s due to its significance in food and feed production [2]. Globally pigeonpea is cultivated on 6.3 million hectares area with production of 5.32 million tons[4]. India is the leading producer contributing 4.34 million despite this, India imports 0.88 million tons of pigeonpea. The productivity of pigeonpea has significantly been impacted by major diseases such as fusarium wilt (FW), sterility mosaic disease (SMD) and phytophthora blight. SMD, caused by pigeonpea sterility mosaic virus (PSMV) corresponding to the emaraviruses, transmitted by the eriophyid mite (*Aceria cajani*) [5]. More than 90% of yield losses have been reported when SMD infects crops at an early stage [6]. In the major pigeonpea growing areas of Indian subcontinent, FW-resistant varieties are available, although many remain susceptible to SMD. For instance, elite cultivars such as Maruti (south India) and JKM189 (central India) are susceptible to SMD, further exacerbating the spread of mites and viruses.

Molecular markers have been successfully used in various genomics-assisted breeding programs in legumes for improving traits such as late leaf spot (LLS) and rust in groundnut [7–9] and fusarium wilt and ascochyta blight in chickpea [7]. Therefore, the development of diagnostic markers is a key to accelerating breeding efforts through marker-assisted breeding. To develop reliable and promising markers, QTL-Seq is one of the most rapid and cost-effective approaches and can be implemented in early (F_2_) as well as in advanced generations (F_8_). This approach was initially developed in rice and used to identify the genomic regions associated with blast resistance and seedling vigor [10]. Subsequently, it is deployed in a variety of crops for multiple traits [11] including groundnut for rust and LLS resistance [12].

Earlier, in pigeonpea, QTL-seq was used to identify genomic regions for FW and SMD resistance in RIL population ICPL20096 × ICPL332 but no success [13]. However, the use of common resistant and susceptible bulks for both FW and SMD might have hindered with QTL detection. The same dataset was then used to identify 26 genome-wide InDels affecting 26 candidate genes [14]. Again, three populations [ICPL20096 × ICPL332, ICPL20097 × ICP8863, and ICP8863 × ICPL87119 (F_2_) population] were used to map genomic regions for SMD resistance using genotyping-by-sequencing (GBS) [15]. Among these, two major QTL regions were identified in ICP8863 × ICPL87119 on chromosome Cc03 (*qSMD3.1*) with 34.3% PVE and LOD 2.8, and on chromosome Cc11 (*qSMD11.3*) with 24.2% PVE and LOD 5.8. Therefore, to further investigate these QTL regions, we advanced ICP8863 x ICPL87119 F_2_ population up to F_8_ generation to develop a RIL population comprised of 153 lines. All these studies were performed using the older version of ‘Asha’ pigeonpea reference genome v1.0, assembled using short reads sequencing [16], therefore the identified genomic regions for SMD resistance have shown less applicability. Interestingly, in this study, we have used the improved telomere-to-telomere ‘Asha’ reference genome v2.0 assembled using Hi-Fi long-read sequencing in combination with Hi-C (High throughput chromosome capture) [17]

In this study, QTL-Seq analysis using improved genome assembly and advanced RIL population (ICP8863 × ICPL87119) identified highly significant (*p* value ≥99% confidence interval) genomic region on chromosome Cc04, harboring non-synonymous variants (SNPs and InDels) and potential disease resistance genes. From this study, KASP markers were designed from identified genomic regions and validated in the entire RIL population (ICP8863 × ICPL87119). Moreover, Indel markers were validated on the 21 resistant and 22 susceptible genotypes of WGRS panel to deploy in genomics-assisted breeding programs to improve SMD resistance.

## 2. Material and methods

### 2.1 Background of the parents and RIL population

In this study, a RIL population comprised of 153 RILs was developed from a cross between susceptible ICP8863 (Maruti) (SMD severity score: 100%) and resistant ICPL87119 (Asha) (SMD severity score: 2%). RIL population and both parents were phenotyped for SMD severity at ICRISAT, Hyderabad, India. The male parent ICPL87119 (Asha), is a medium-duration, high-yielding, and resistant to FW and SMD. While the ICP8863 (Maruti), a popular FW resistant cultivar but susceptible to SMD was used as a female parent.

### 2.2 Phenotyping for the sterility mosaic disease (SMD) and construction of bulks

RIL population along with parents were planted in sick plot specially designed for SMD screening at ICRISAT, Hyderabad (17.5111° N, 7832752° S) during rainy 2023. 15 days old plants were inoculated at the two-leaf stage using the leaf stapling technique [18]. SMD incidence was recorded at three growth stages in one particular season. The first observation was recorded for 30 days post-inoculation (DPI), followed by the second observation at 60 DPI and third was recorded at the flowering stage. Disease incidence scale ranging from 0% (highly resistant) to 100% (highly susceptible) was used to categorize disease incidence into four groups: 0-10% (resistant), 11-20% (moderately susceptible), 21-40% (susceptible), >40% (highly susceptible). The overall disease incidence was calculated using the below-mentioned formula [18]:

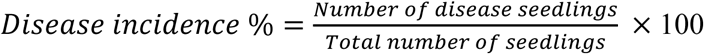

Based on the phenotyping data for SMD disease incidence (%) generated on the RIL population, a total of 18 resistant and 18 susceptible RILs were used to construct resistant (R-bulk) and susceptible (S-bulk), respectively (Supplementary table1, Figure 1a).

**Figure 1:**
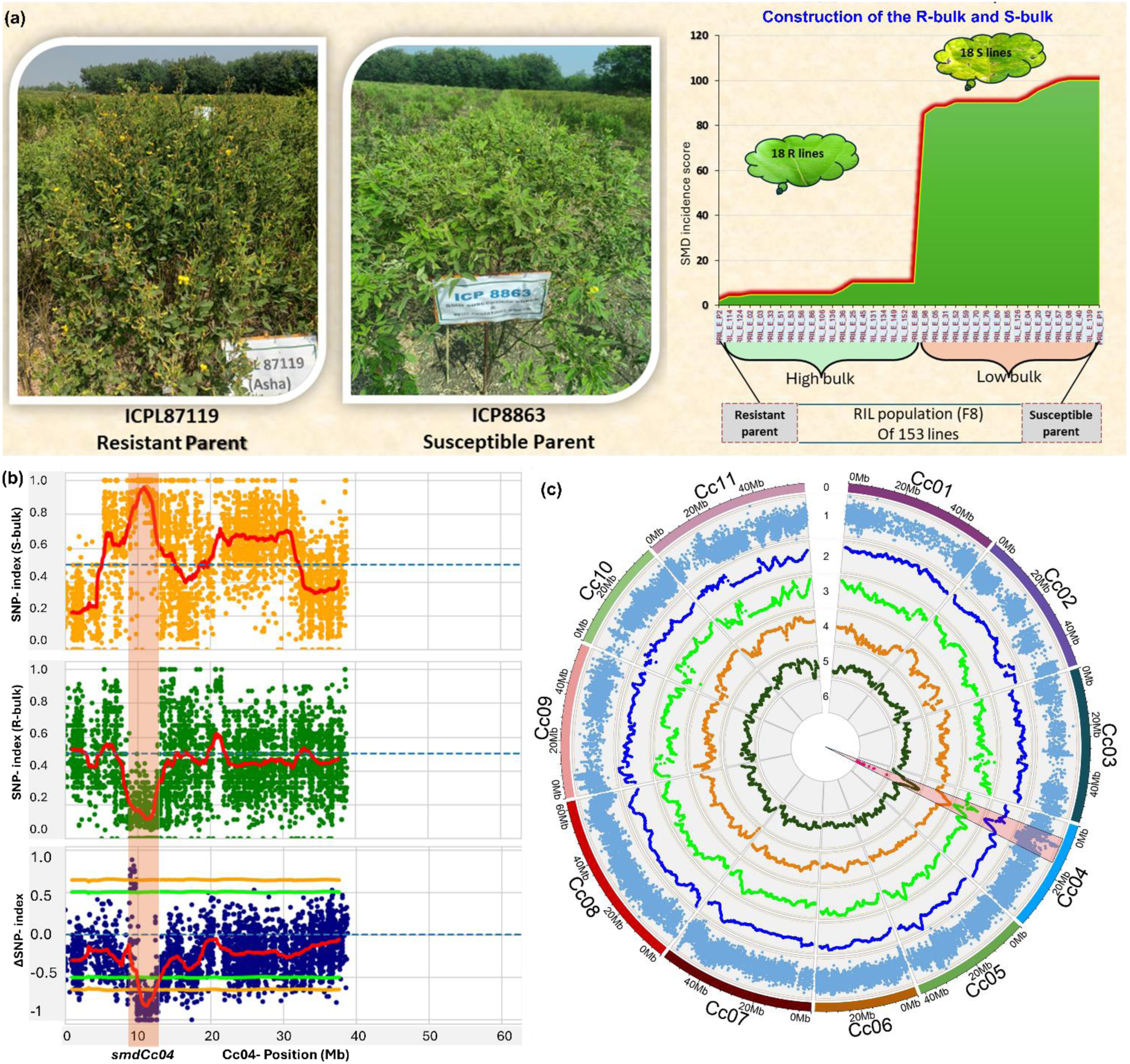
QTLseq analysis and identification of associated genomic region on chromosome Cc04 for SMD **a**. Phenotypes of resistant and susceptible parents and construction of resistant or R-bulk and susceptible or S-bulk by selecting extremely resistant and susceptible RILs for sterility mosaic disease from RIL population ICP8863 × ICPL87119. **b.** SNP-index plot for low bulk in orange color, high bulk in green color, and Δ-SNP index plot in blue illustrates the genomic region *smdCc04* (highlighted in light orange color). **c.** Circos plot illustrates genome-wide variants identified for SMD resistance in pigeonpea. The tracks go from outside to inside. 0. 11 chromosomes of pigeonpea 1. Genome-wide variants identified on each chromosome. 2. Δ-SNP index (P > 0.01) using resistant parent ICPL87119. 3. Δ-SNP index (P > 0.01) of susceptible parent ICPL88863. 4. SNP index for S-bulk. 5. SNP index for R-bulk. 6. Non-synonymous SNPs identified on chromosome Cc04 and Cc11.

### 2.3 DNA extraction, library construction and sequencing of bulks and parents

High quality DNA was extracted from each line in the RIL population including the parents using the NucleoSpin^®^ 96 Plant II (Macherey-Nagel) Kit, and DNA quality was checked on a 0.8% agarose gel and quantity was estimated using Nanodrop spectrophotometer (Thermofisher Scientific, USA) as explained in [19] Based on the SMD disease incidence (%) on 153 RILs, the DNA of 18 resistant and 18 susceptible RILs was pooled to construct R-bulk and S-bulk respectively (Figure 1a) as detailed in [12]. Quality of pooled DNA for both bulks was checked on a 0.8% agarose gel before library preparation. Further, a single library was prepared for each sample using TruSeq DNA Sample Prep kit, and libraries of two bulks and susceptible parent (ICP8863), the quality of the libraries was checked using Qubit fluorometer and sequenced using the Illumina HiSeq 2500 platform. The sequencing data for resistant parent was already available.

### 2.4 Development of reference guided assembly and bi-directional QTL-Seq analysis

The resistant parent ICPL87119 ‘Asha’ used in this study is already being used for developing reference genome assembly in pigeonpea [16]. Therefore, the susceptible parent, ICP8863 was aligned with the improved reference genome assembly [17] to identify the variants between the parents. The reads of R-bulk and S-bulk were aligned with both parents, and variants were called for both bulks to calculate SNP-index (Table 1, Supplementary figure 1, Supplementary figure 1). The statistics of generated sequencing reads were estimated using the raspberry tool of NGS-QCbox [20].

**Table 1.** Details of sequencing data generated for both bulks and parents.

| Sample | Pedigree | Phenotyping<br>of SMD (%) | Illumina sequencing |  |  |  | Data processing |  |  |
| --- | --- | --- | --- | --- | --- | --- | --- | --- | --- |
|  |  |  | Data<br>generated<br>bases (Gb) | Reads<br>(Million<br>reads) | Read<br>depth<br>(X) | Genome<br>coverage<br>(%) | Trim<br>bases<br>(Gb) | Trim<br>reads<br>(Million<br>reads) | Alignment<br>(%) |
| Parents |  |  |  |  |  |  |  |  |  |
| ICP8863 | Selection from<br>landrace | 100 | 12.42 | 82.27 | 22.35 | 92.5 | 11.67 | 78.06 | 98.35 |
| ICPL87119 | C11 9 ICP1-6-W3-W | 2 | 25.14 | 167.58 | 45.33 | 91.75 | 23.77 | 160.93 | 98.21 |
| Mapping population |  |  |  |  |  |  |  |  |  |
| R-Bulk | ICP8863 × ICPL87119 | 0-10 | 13.61 | 90.11 | 24.42 | 93.12 | 86.14 | 86.14 | 98.11 |
| S-Bulk | ICP8863 × ICPL87119 | 85-100 | 15 | 99.33 | 26.9 | 93.2 | 94.35 | 94.35 | 98.08 |
| Total |  |  | 91.31 | 606.87 | 164.33 | 462.32 | 239.7 | 580.41 | 490.96 |

**ICP8863;** Susceptible parent, **ICPL87119:** Resistant parent, **R-Bulk:** Resistant bulk, **S-Bulk:** Susceptible bulk.

QTL-seq analysis was performed by using QTL-seq pipeline developed by [10]. All the SNPs and InDels above the 99% confidence interval (CI) and 95% CI from the significant region (peak region) were extracted as guided by the sliding windows (0.2 Mb). A SNP variation index for each SNP position was calculated for resistant and susceptible bulks by using the following formula [10]:

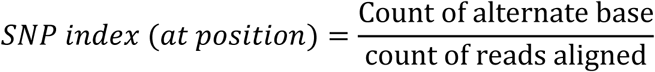

SNPs with homozygous alleles in susceptible and resistant bulks, were used for the calculation of the Δ-SNP index. For the calculation of Δ-SNP index, the SNP index of S-bulk was subtracted from the SNP index of the R-bulk (Supplementary Figure 3). The SNPs having Δ-SNP index = - 1 and near to-1, were considered as favorable alleles, contributed by resistant parent. While the alleles with Δ-SNP index = 1 and near to 1, were considered as non-favorable alleles contributed by susceptible parent. Among these SNPs and InDels, the synonymous and non-synonymous variants were investigated using snpeff version 5.2c [21] and snpsift version 5.2c [22]. All the genes were extracted from the peak region and their functions were identified by using the information available in the annotation file and by using the protein sequences of the respected genes. Preferentially, the function of candidate genes having nonsynonymous variants were identified and their relationship with disease resistance was also identified (Table 2). Circos plot illustrating the SNP index and Δ-SNP index calculated from both the parents and bulks was developed using ShinyCircoseV2.0 (https://venyao.xyz/shinycircos/) [23] (Figure 1c).

**Table 2.**
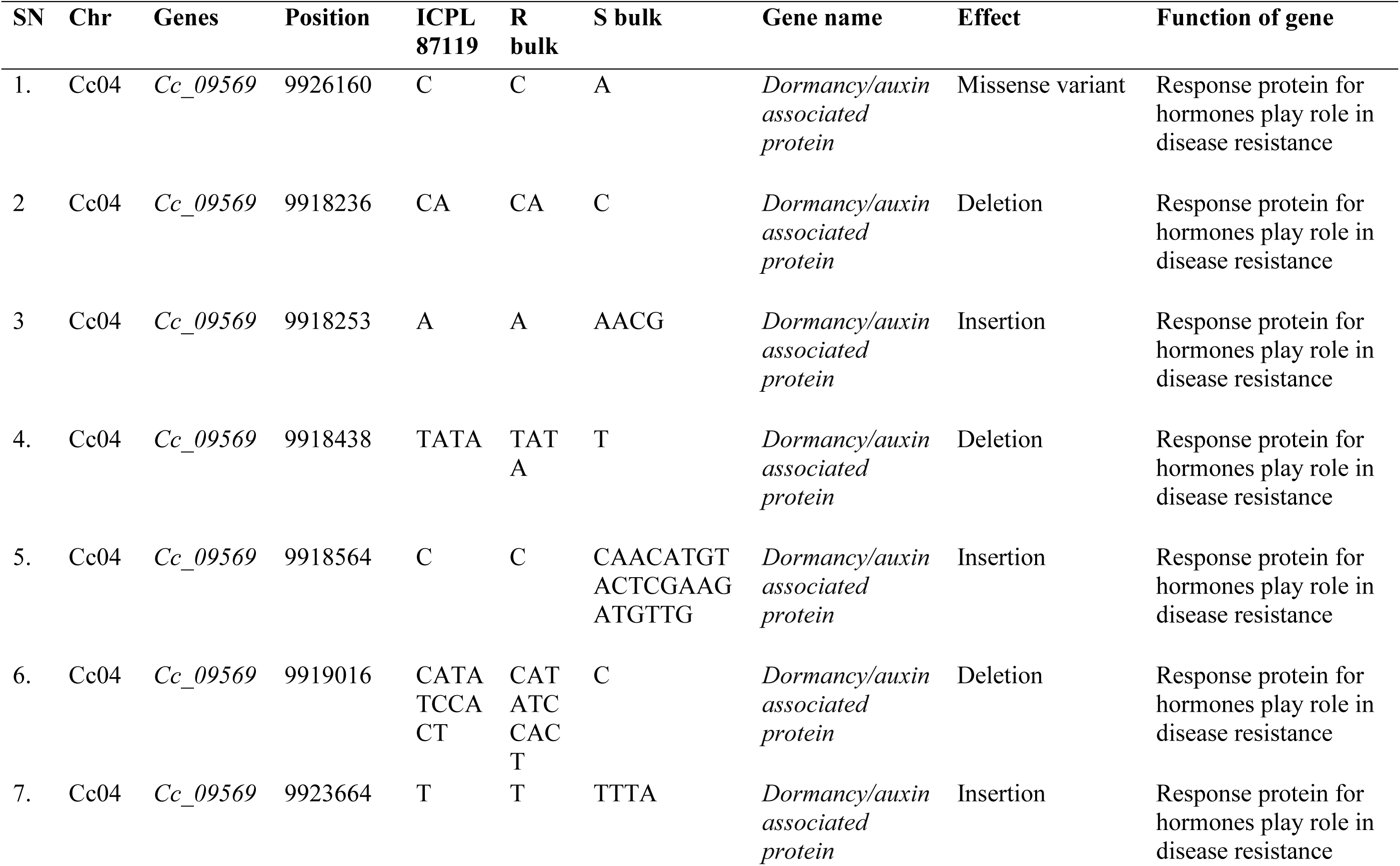
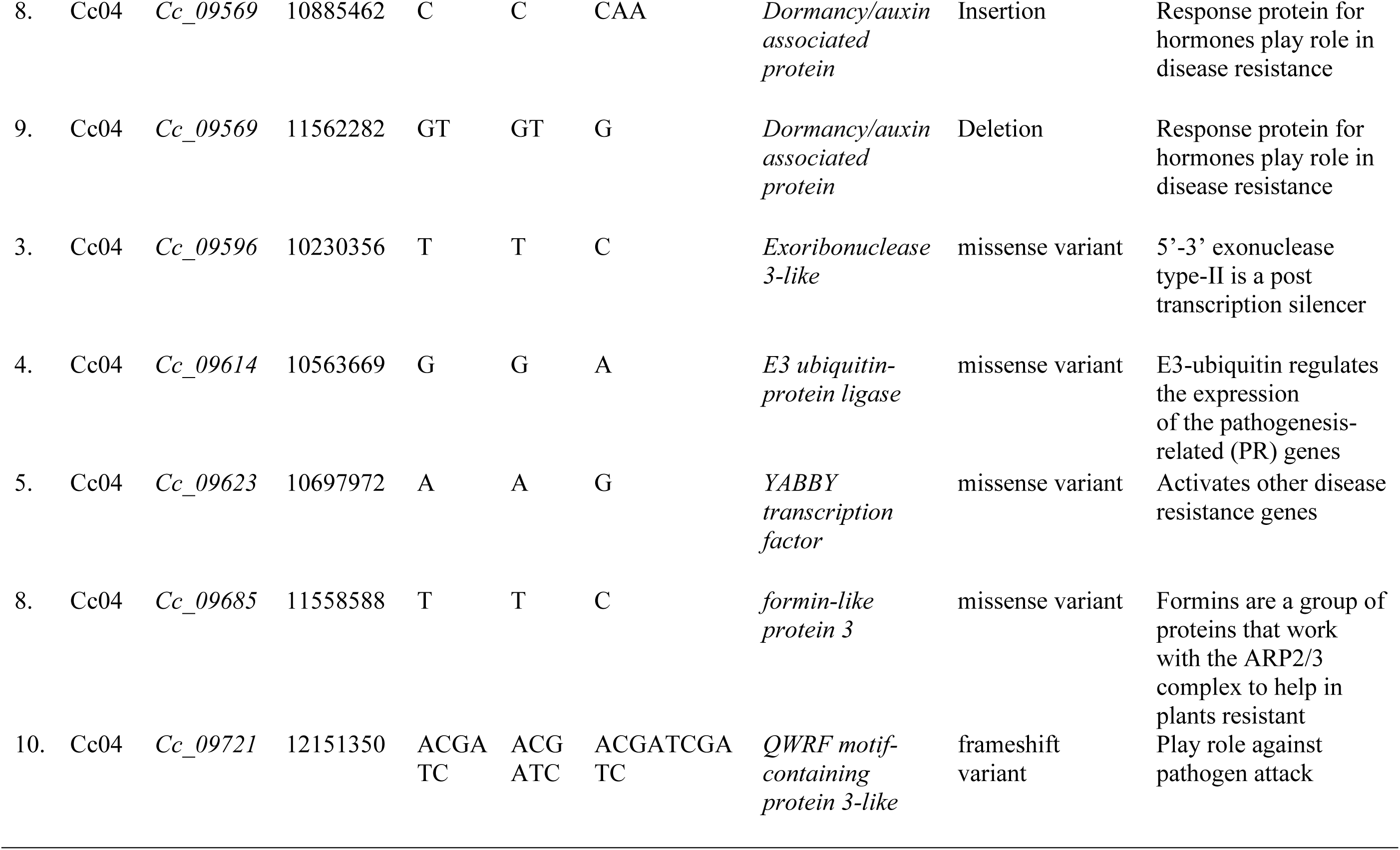
List of nonsynonymous/Indel variants and corresponding candidate genes identified for SMD resistance.

### 2.5 Allele mining for validation of candidate genes in the identified QTL genomic region

The markers from the genomic region (*smdCc04*) on chromosome Cc04 were mined for favorable alleles in SMD resistant and susceptible lines using whole genome resequencing (WGRS) data (Supplementary table 2). A total of 4 insertions (8, 21, 3, and 3 bp), 2 deletions (1 and 9 bp), and 8 SNPs from *smdCc04* region were used for allele mining in resistant and susceptible lines. The functional validation of the identified Indels was done by identifying regulatory elements using Plantpan4.0 webtool (https://plantpan.itps.ncku.edu.tw/).

## 3. Results

### 3.1 Phenotypic variability for sterility mosaic disease in RIL population

SMD disease incidence (%) for the RIL population ranged between 5% to 100% with a mean of 47.3%. Based on SMD disease incidence (%), 18 susceptible and 18 resistant RILs were selected from the 153 RILs for the construction of susceptible and resistant bulks. The phenotypic score for the resistant bulk ranged from 5% to 10% with a mean of 6.6%, and for susceptible bulks 85% to 100% with a mean of 88% (Figure 1; Supplementary table 1).

### 3.2 Whole genome sequencing and variant calling

A total of 13.61 Gb data was generated (92.5% coverage) for the R-bulk with 90.11 million reads at a read depth of 24.42X. For S-bulk, 15 Gb data was generated (91.75% coverage) with 99.33 million reads at a depth of 26.9X. For susceptible parent ICP8863, 12.42 Gb data was generated (92.5% coverage) with 82.27 million reads at a read depth of 22.35X. The resistant parent ICPL87119 (Asha) was sequenced earlier with coverage (91.75%) [17]. The trimmed and filtered high-quality reads of the R-bulk (86.14 million reads) and S-bulk (94.35 million reads) mapped against the newly developed reference guided assembly of ICPL87119. The alignment percentage of R-bulk was 98.11% followed by S-bulk 98.08%. At a read depth ≥ 7 in both the bulks and ≥0.3 SNP index in either of the bulks were considered as a selection criterion for resistant loci (Table 1).

A total of 603,135 genome-wide variants were identified, with 137,019 heterozygous and 465,916 homozygous variants between R-bulk and S-bulk. A total of 29,780 polymorphic SNPs and 9,256 polymorphic InDels recovered after aligning the high and low bulks with the reference-guided assembly. Minimum 1581 SNPs were obtained on chromosome Cc01 and maximum 4975 SNPs on chromosome Cc08. Similarly, a minimum of 520 InDels were obtained on chromosome Cc03 and maximum 1346 InDels were obtained on chromosome Cc08 (Supplementary table 3).

### 3.3 Identification of genomic regions and candidate genes for sterility mosaic disease

QTL-seq analysis identified one potential genomic region on chromosomes Cc04 (*smdCc04*). On chromosome Cc04, the genomic region *smdCc04* of 3.2 Mb (9.3-12.5 Mb) contained 3,540 variants including 2,324 SNPs, and 1,103 InDels. Among these variants, 266 (189 SNPs and 77 InDels) were identified at the 95% confidence interval (CI) (*p < 0.05*), and 229 (173 SNPs and 56 InDels) were identified at 99% CI (*p < 0.01*) (Figure 1c, Supplementary table 4).

The identified genomic region was mined for candidate gene discovery using Asha genome v2.0. The genome variants such as non-synonymous, intron splicing region variants, and premature start codon variants corresponding genes were referred to as candidate genes. Especially, non-synonymous variants such as missense, frameshift mutations, stop-gained, stop-lost, start gain, and start-lost variants were considered as causative variants impacting the function of candidate genes (Supplementary table 5).

In the genomic region *smdCc04,* on chromosome Cc04, a total of 211 genes were identified, of which 28 are involved in the pathway of SMD resistance mechanism (Table 3; Supplementary table 6). In this region, missense variants were corresponding to important candidate genes including *Dormancy/auxin associated protein* (*Cc_09569*), *exoribonuclease 3-like* (*Cc_09596*), *E3 ubiquitin-protein ligase* (*Cc_09614*), *YABBY transcription factor* (*Cc_09623*), *formin-like protein 3* (*Cc_09685*), *QWRF motif-containing protein 3-like* (*Cc_09721*). Splice-region-variants were found in probable *CCR4-associated factor* (*Cc_09591*), *mitogen-activated protein kinase A-like* (*Cc_09636*), and *aconitate hydratase* (*Cc_09636*) genes. Interestingly, six frameshift variants and one missense were identified in the region of *Dormancy/auxin associated protein* (*Cc_09569*) gene which is one of the signal receptor genes (Table 2, Figure 2). This gene has seven significant variants, six are Indels and one is missense SNP variation.

**Figure 2:**
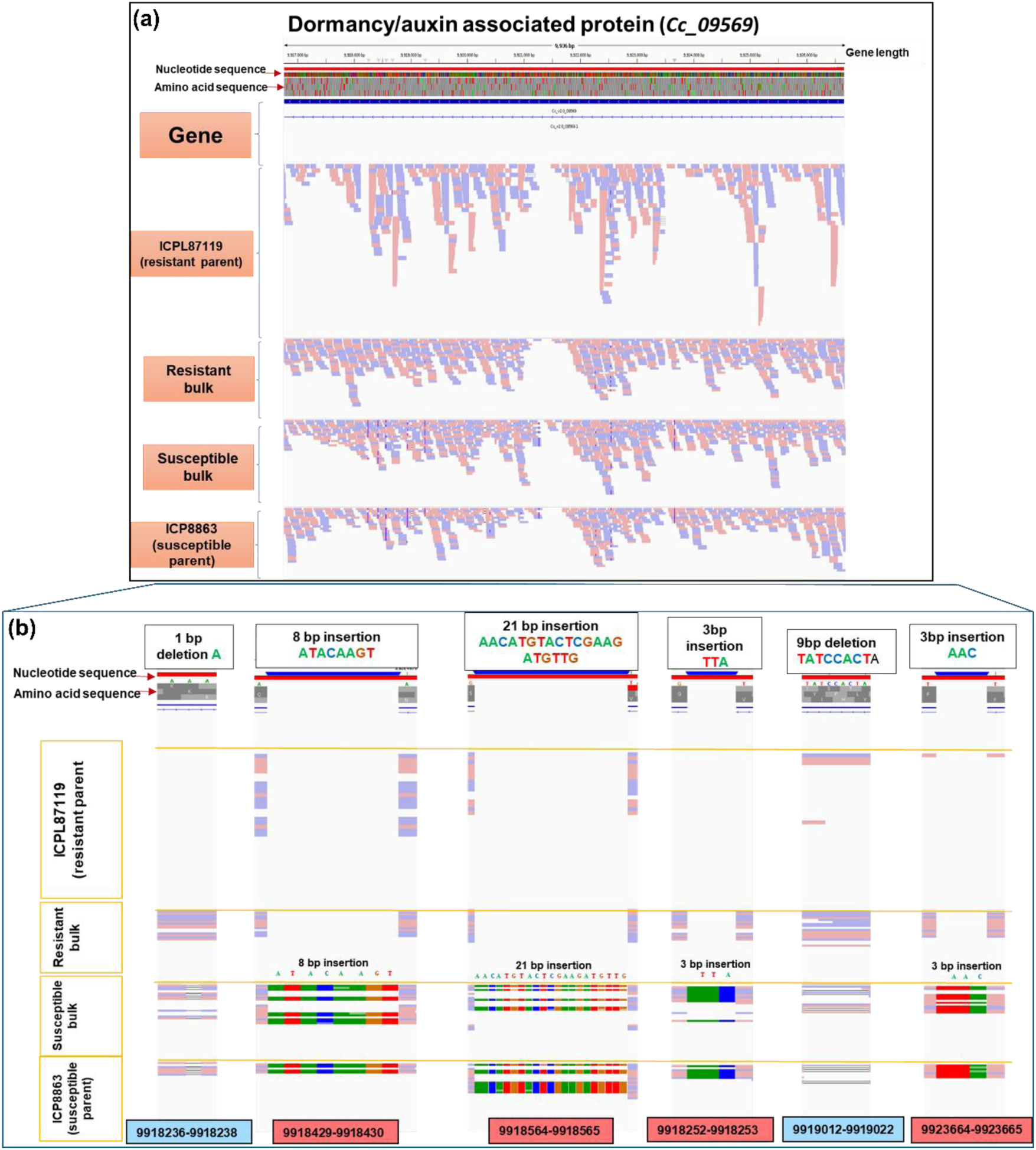
Insertion and deletions identified in associated genomic region *smdCc04* for SMD resistance **a**. Sequencing data of resistant and susceptible parents with their respective bulks aligned to genomic region of Cc_09569 pigeonpea genome version 2. Sequence represents nucleotide sequence of genes and amino acid sequence of respected protein. Red stars are for stop codon; green color is for methionine. **b.** Represents insertion and deletion for the gene Cc_09569 in resistant and susceptible parent with their respective bulks. Sequence represents nucleotide sequence of gene and amino acid sequence of respected protein.

**Figure 3:**
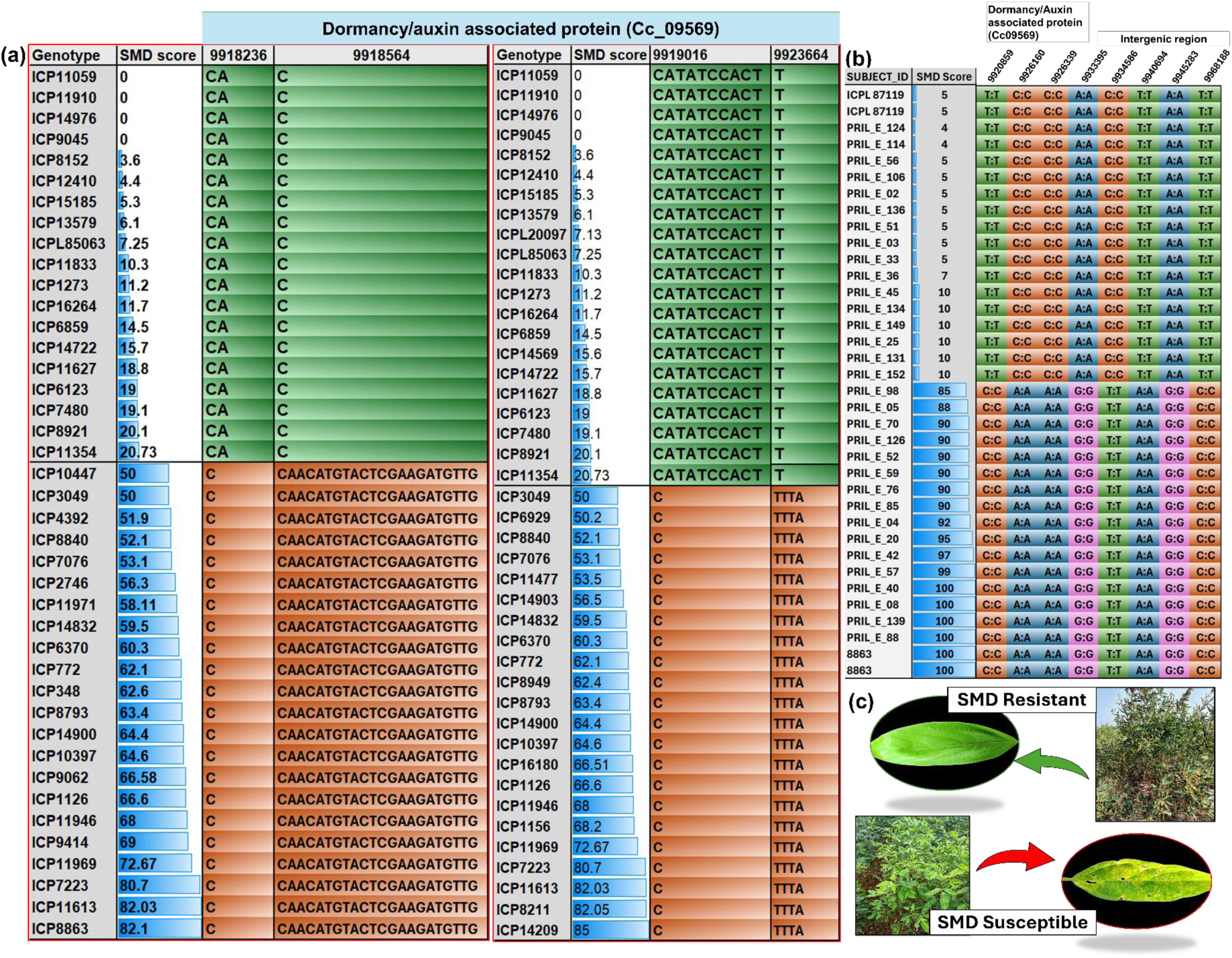
Validation of Indels and missense SNP present in Dormancy/auxin associated protein (Cc_09569) from *smdCc04* (9.3-12.5 Mb) region **a.** Allele-mining of the Cc04 markers on the contrasting lines of SMD trait. From *Dormancy/auxin associated protein* (Cc_09569) one deletion and one insertion marker validated on a one set of 19 resistant and 22 susceptible genotypes, and one insertion and one deletion marker validated on another set of 21 resistant and 22 susceptible genotypes **b.** Validation of KASP markers. From *Dormancy/auxin associated protein* (Cc_09569) three intronic SNP validated on 16 resistant and 16 susceptible lines of ICP8863 × ICPL87119 population. The other five validated markers are housed in the intergenic region. **c.** Phenotype of SMD resistant and SMD susceptible.

**Table 3:**
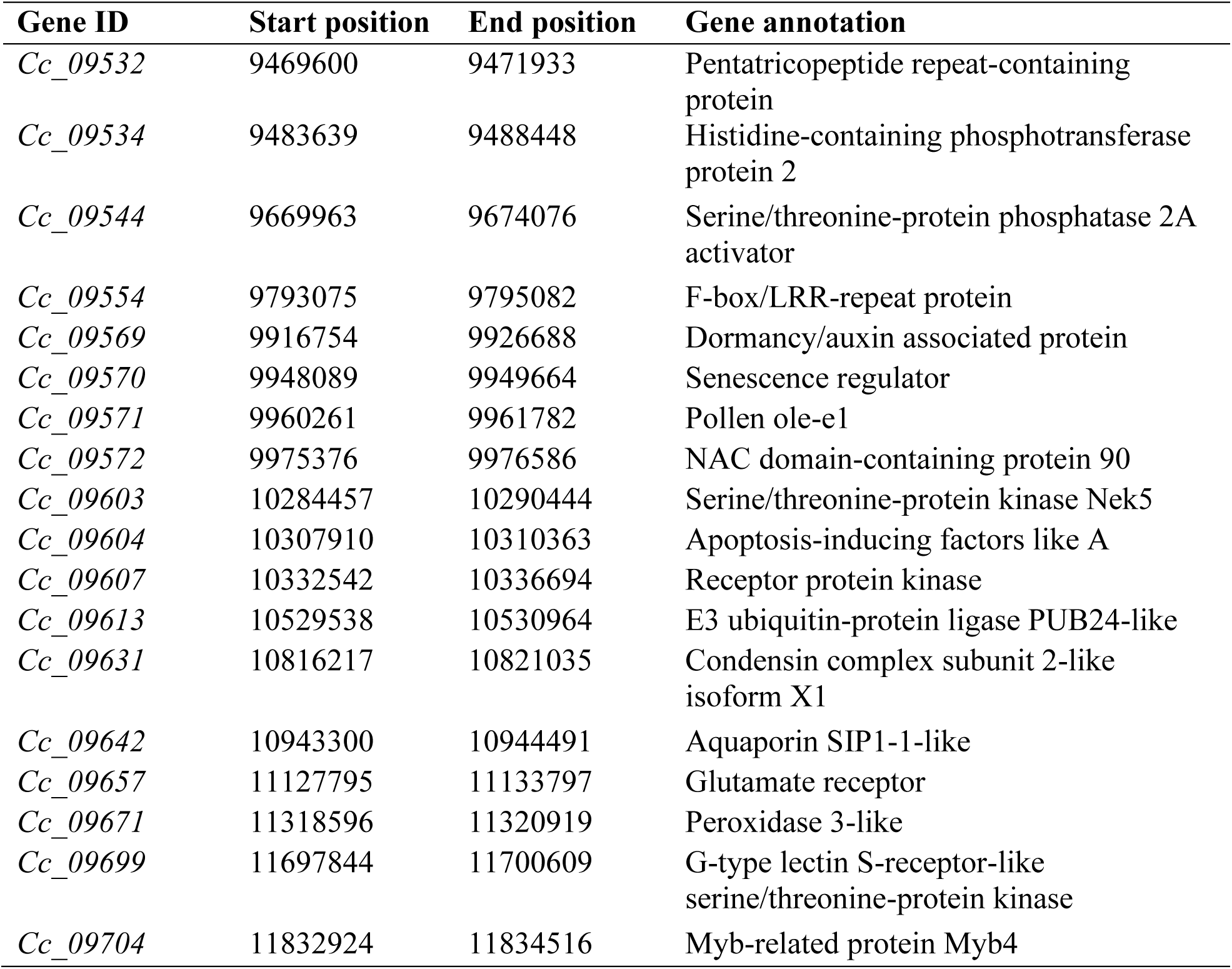
List of disease resistance related genes from *smdCc_04* SMD genomic regions.

### 3.4 Identification of SNPs and mining favorable alleles in whole genome re-sequencing data of resistant and susceptible genotypes

Allele-mining of putative markers from the *smdCc04* region was performed in the whole genome-resequencing data of resistant and susceptible genotypes. Alleles from the resistant and susceptible parents were mined in the 19 resistant and 22 susceptible genotypes for Cc04_9918236 (1bp deletion) and Cc04_9918564 (21bp insertion) markers. For marker Cc04_9919016 (9bp insertion) and Cc04_9923664 (3bp insertion) alleles from the resistant and susceptible parents were mined in the 21 resistant and 22 susceptible genotypes. Moreover, for one missense SNP marker Cc04_9926160 allele mining was done in 5 resistant and 5 susceptible genotypes. All these markers were corresponding to *dormancy/auxin associated protein* (Cc_09569) gene. Auxin significantly influences plant-microbe interactions, particularly between plant hosts and disease-causing pathogenic microorganisms (Figure 4).

**Figure 4:**
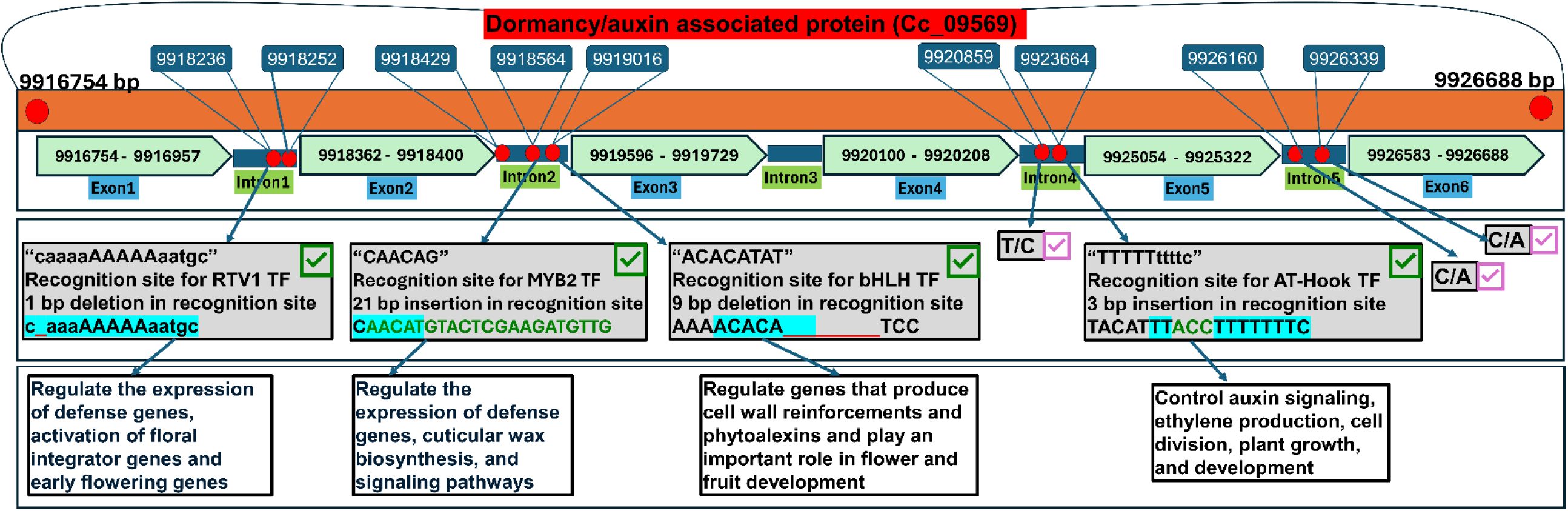
Four functionally validated Indel markers (mentioned by green check mark) housed in the intronic region of the dormancy/auxin associated protein (Cc_09569). Cc04_9918236 is identified in Intron1, Cc04_9918564 and Cc04_9919016 in Intron2, and Cc04_9923664 is in Intron4. Marker Cc04_9918236 disrupted the transcription factor recognition site (caaaaAAAAAaatgc) for RTV1 transcription factor, marker Cc04_9918564 disrupted the transcription factor site (CAACAG) for MYB2 transcription factor, marker Cc04_9919016 disrupted the transcription factor site (ACACATAT) for bHLH transcription factor, and marker Cc04_9923664 disrupted the transcription factor site (TTTTTttttc) for the AT-hook. Moreover, three validated SNP markers (mentioned by purple check mark) housed in the intronic region of dormancy/auxin associated protein (Cc_09569). Cc04_9920859 is identified in Intron4, Cc04_9926160 and Cc04_9926339 are identified in Intron5.

### 3.5 Validation of the InDel and SNP markers and their functional validation

We have identified a total of six InDel markers in *dormancy/auxin associated protein* (Cc_09569), namely, Cc04_9918236, Cc04_9918564, Cc04_9919016, Cc04_9923664, Cc04_9918429, and Cc04_9918252 (Figure 2). Among these, 4 are InDel markers validated in WGRS sequencing panel, namely, Cc04_9918236, Cc04_9918564, Cc04_9919016, Cc04_9923664. These four functional markers are housed in the intronic region of the *dormancy/auxin associated protein* (Cc_09569). Cc04_9918236 is identified in Intron1, Cc04_9918564 and Cc04_9919016 in Intron2, and Cc04_9923664 is in Intron4. Marker Cc04_9918236 disrupted the transcription factor recognition site (caaaaAAAAAaatgc) for RTV1 transcription factor, marker Cc04_9918564 disrupted the transcription factor site (CAACAG) for MYB2 transcription factor, marker Cc04_9919016 disrupted the transcription factor site (ACACATAT) for bHLH transcription factor, and marker Cc04_9923664 disrupted the transcription factor site (TTTTTttttc) for the AT-hook (Figure 4a).

We have identified and sent 15 SNP markers for KASP validation. Among 15 SNPs, a total of 8 markers were validated on the validation panel. Three SNP markers (Cc04_ 9920859, Cc04_ 9926160, and Cc04_ 9926339) are housed in the *dormancy/auxin associated protein* (Cc_09569). While rest of the five SNP markers (Cc04_9933395, Cc04_ 9934586, Cc04_ 9940694, Cc04_ 9945283, and Cc04_ 9968188) are housed in the intergenic region between *dormancy/auxin associated protein* (Cc_09569), senescence regulator (Cc_09570), and protein ole e-1 (Cc_09571). A total of 151 lines (out of 153 lines) of the ICP8863 × ICPL87119 RIL population (PRILE) were sent for validation. A total of 69 PRIL resistant lines (out of 77 resistant lines) have been validated with the resistant allele, and 58 susceptible lines (out of 75 susceptible lines) have been validated with the susceptible allele. 11 lines have shown heterozygous alleles (Figure 5 and Supplementary table 10). Resistant bulk and susceptible used in this study contain 18 resistant and 18 susceptible lines, respectively. Out of 18 resistant lines, 16 have shown resistant alleles, while out of 18 susceptible lines, 16 have shown susceptible alleles (Figure 4.b).

**Figure 5:**
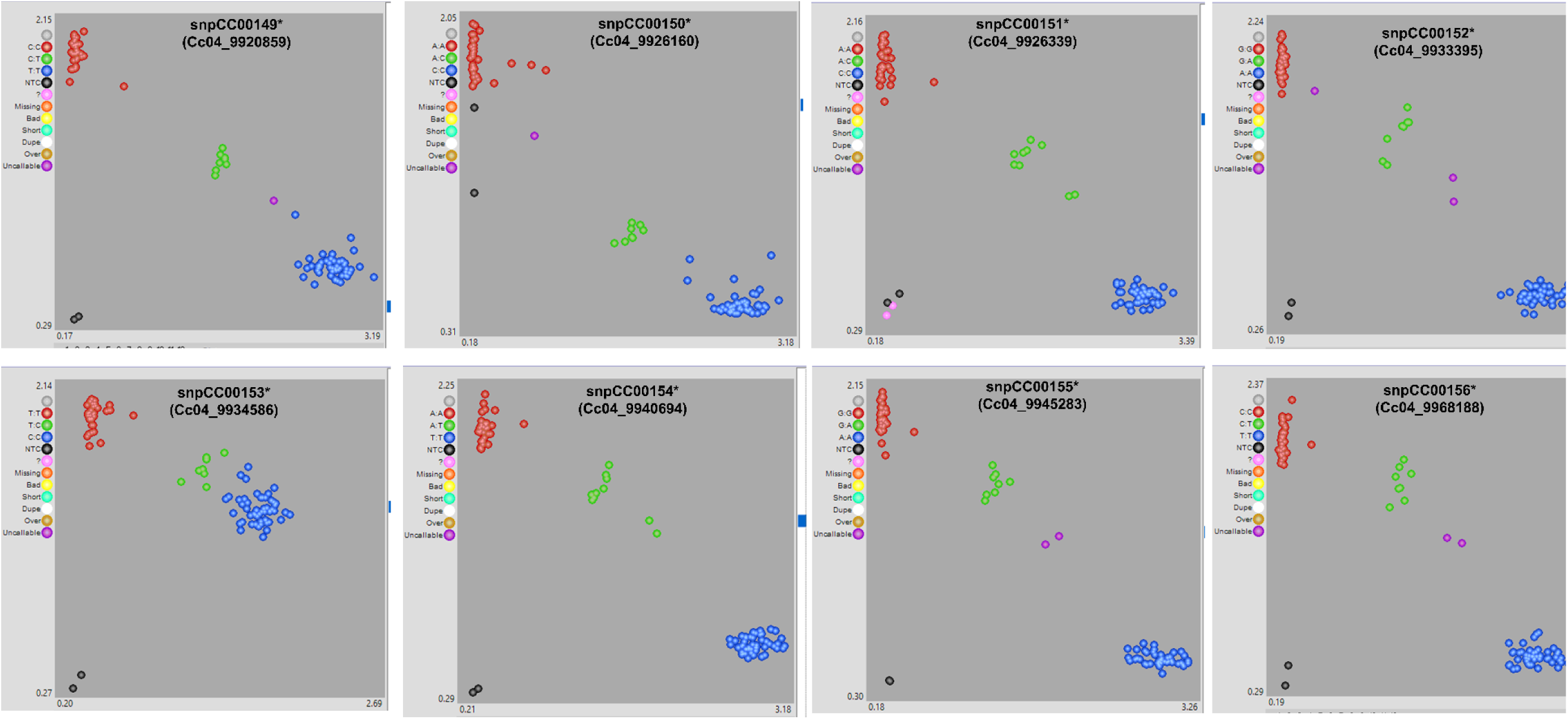
The pictorial representation of the KASP assays conducted on a RIL population (ICP8863 × ICPL87119) of 140 lines including parents. Red color indicates the favorable allele, blue color indicates the non-favorable and green color indicates the heterozygotes. Star indicates Intertek IDs.

## 4. Discussions

Pigeonpea is more an orphan crop and has now ample genomic resources for gene discovery and genomic-assisted breeding (GAB). SMD is a devastating disease that results in a 95-100% yield penalty in the Indian sub-continent. It is essential to identify associated genome regions resistant genes from various resistance sources and introgress them into elite cultivars to mitigate resulting yield losses due to SMD. The whole genome sequencing and re-sequencing approaches have proven to be reliable and effective for high-resolution trait mapping [24,25]. In the past decade user-friendly bioinformatics pipelines such as QTL-Seq, Mut-Map, Indel-Seq have increased the accuracy of trait mapping and marker discovery for complex traits in various crops such as rice [10], pigeonpea [13,26] and groundnut [12,27,28]. The QTL-seq approach is resource efficient because it requires sequencing of fewer samples, a time-efficient method that yields high-quality results for the targeted oligogenic trait.

Earlier, QTL-seq on ICPL20096 × ICPL332 RIL population was used in common resistant and susceptible bulks for FW and SMD to identify genomic regions for FW and SMD using an old version of genome Asha developed using short reads [13]. No QTL could be detected maybe because common bulks were made for both the traits, and the phenotypic effect of QTL was minor/less. QTL-Seq study on ICP8863 × ICPL87119 RIL population identified one genomic region associated with SMD resistance on chromosomes Cc04. We selected this RIL population because major QTLs detected in F2 mapping population of our previous study [15]. Further, we used the improved genome assembly of ‘Asha’ as a reference genome [17] using RIL population (ICP8863 × ICPL87119, F_8_ generation). This RIL population was advanced using the F_2_ mapping population previously utilized in the GBS-based QTL identification study for SMD resistance [15]. The initial genome assembly, covering 40.86% of the genome [16], was recently upgraded to achieve 91.35% genome recovery [17], enhancing the reliability of sequencing-based approaches.

A region on Cc04 (*smdCc04*), spanning 9.3-12.5Mb, contains variants above the 99% confidence interval (CI) and 12 non-synonymous variants (6 SNPs and 6 Indel) impacting the function of six disease resistance-related genes. Among 12 variants, one missense SNP and six Indels variants were identified in the *Dormancy/auxin associated protein* gene (*Cc_09569*). This protein is generally responsive to hormones (Such as auxin) involve in biotic defense response [29]. A missense variant (Cc04_10563669) was identified in *E3-ubiquitin protein ligase* (*Cc_09614*), which plays a critical role in the expression of pathogenesis-related (PR) genes during plant disease resistance from pathogen perception to execution [30,31]. A missense variant at (Cc04_10697972) was identified in *YABBY-protein* (*Cc_09623*). The *YABBY transcription* family interacts with other disease-resistance genes like *CRR3*, playing a crucial role in plant disease resistance [32]. In pineapple plants, the *YABBY transcription factor* triggers the accumulation of ABA and ethylene in response to pathogen attack [33]. A splice region variant (Cc04_10885462) was identified in the region of mitogen-activated protein kinase A-like (MAPK) (Cc_09636). MAPKs play a crucial role in signaling plant defense against pathogens and microbes by developing MAPK cascade. For instance, in cotton *MPK9*, *MPK13*, and *MPK25* are upregulated when inoculated with Verticillium wilt causing *Verticillium dahliae*. Silencing these genes makes the plant more susceptible to *V. dahlia* [34][34]. The MAK3/MPK6 cascade and calcium signaling pathway, activated during plant immunity, converge on *WRKY33* to regulate the biosynthesis of camalexin, a crucial phytoalexin for disease resistance in Arabidopsis [35].

Interestingly, four Indels and (Cc04_9918236, Cc04_9918564, Cc04_9919016, and Cc04_9923664) were identified in the *Dormancy/auxin associated protein (Cc_ARP)* (*Cc_09569*). These mutations are responsible for the disruption of *Cc_ARP* leading to the hinderance of auxin signaling pathways. The downregulation of *Dormancy/auxin associated protein* enhances the plant susceptibility to infection by *Tobacco mosaic virus* (TMV), *Pectobacterium carotovorum subsp. carotovora*, and *Phytophthora parasitica var*. *nicotianae*. Similarly, the knockdown of *TaARP* gene resulted in higher susceptibility to fungus *Blumeria graminis f. sp tritici* in *Triticum aestivum* [36,37]. *Dormancy/auxin associated protein* gene is responsive to the signaling hormones involved in plant disease resistance, growth, and development, such as salicylic acid (SA), jasmonic acid (JA), ethylene, and auxins [29]. It is reported that the activation of auxin signaling enhances the resistance against plant viruses [38]. The overexpression of auxin signaling repressor in lines of rice showed susceptible response to the *Rice black streaked dwarf virus* (RBSDV) [39]. This study also revealed that the jasmonic acid pathway may also be influenced by the auxin auxin-signaling mediated response to the RBSDV in rice. The ARF factor was targeted by several viral proteins and in each case, it is found that these interactions benefited for the viral infections. This suggests that manipulation in auxin signaling is the common pathogenicity strategy in plant viruses [40].

Hormones play a most crucial role in the development of reproductive tissues by initiating and regulating the growth and maturation of reproductive organs and controlling the timing of important reproductive events [41]. The proper signaling of these hormones is mediated by response proteins such as *Dormancy/auxin associated protein* [42]. Therefore, the sterility in the reproductive tissues of pigeonpea might be due to the loss in the function of *Dormancy/auxin associated protein* gene. *Dormancy/auxin associated protein* (Cc_09569) has 10 variants. Four Indels and three SNP are validated on WGRS sequencing panel and through KASP genotyping respectively. The gene family of this gene shows the important link between response to pathogen and plant growth and development. Therefore, this gene could be considered as the most important in this region and further transcriptomics, proteomics, and metabolomics studies may be performed to explore the actual mechanism in the pigeonpea resistant and susceptible genotypes. In summary, genomic regions *smdCc04* identified for SMD resistance uncovered important disease resistance genes and markers for validation. This study indicated that SMD susceptibility might be due to the disruption of *Dormancy/auxin associated protein* gene (*Ccsmd04*) under infection sterility mosaic disease (SMD).

Identification and validation of the functional markers is the major outcome of any study done by using genomic resources. Indels have potential to disturb the functional associated gene. In our study, we have identified 4 functional InDel markers, namely, Cc04_9918236, Cc04_9918564, Cc04_9919016, Cc04_9923664. These markers are housed in the intronic region of the *dormancy/auxin associated protein* (Cc_09569). Cc04_9918236 is identified in Intron1, Cc04_9918564 and Cc04_9919016 in Intron2, and Cc04_9923664 is in Intron4. Marker Cc04_9918236 have disrupted the transcription factor recognition site (caaaaAAAAAaatgc) for RTV1 transcription factor, marker Cc04_9918564 have disrupted the transcription factor site (CAACAG) for MYB2 transcription factor, marker Cc04_9919016 have disrupted the transcription factor site (ACACATAT) for Basic helix-loop-helix (bHLH) transcription factor, and marker Cc04_9923664 have disrupted the transcription factor site (TTTTTttttc) for the AT-hook. RTV1 transcription factor is a DNA binding domain (B3 domain family) generally controls the flowering time [43]. MYB2 transcription factor regulates the expression of genes that are involved in pathogens and pests’ resistance pathways. This TF also facilitates the biosynthesis of lignin, flavonoids, and cuticular wax. Moreover, this TF regulates the signaling of plant hormones involved in plant disease resistance [44]. The bHLH transcription factors fight plant pathogen by sensing H_2_O_2_, lignin and phytoalexin biosynthesis [45]. AT-hook motif nuclear localized (AHL) family plays an important role in plant disease resistance by regulating the expression of defense-related genes [46].

The expression of the gene is controlled through the binding of transcription factors (TFs) to regulatory genomic regions [47]. Some of the studies report that the presence of the transcription factor recognition site in the intronic region of the gene could play an important role in the gene expression regulation by influencing splicing mechanisms and potentially modifying the rate of transcription elongation within the body of gene. These sites can act as regulatory elements within the intron to control the transcription of the gene and their processing into mature mRNA [48]. Few earlier studies have confirmed that the conserved transcription factor binding sites are required for the appropriate expression of the genes in *Caenorhabditis elegance* [49–52]. In *Arabidopsis thaliana*, the second intron of AGAMOUS (AG) gene contains cis-acting DNA sequences that bind the transcription factors LEAFY and WUSCHEL which control floral fate and floral-specific expression of AG, respectively [53]. Such types of introns that act as a co and post-transcriptional regulation are called as intron-mediated enhancement (IME). Earlier, this phenomenon was reported in maize genes *Alcohol dehydrogenase1* and *Bronze1* [54], and *Shrunken1* [55]. In *Arabidopsis thaliana* it was reported in gene *Phosphoribosylanthranilate transferase1* [56].

## 5. Conclusion

This study identified candidate genes and markers (SNPs and Indels) for SMD resistance in pigeonpea. Most of the genes identified in this study play an important role in Plant-pathogen interaction, growth, development, and signaling. The identified four frameshift mutations (InDel) in *Dormancy/auxin associated protein* gene showed 1 bp insertion, 21 bp insertion, 9 bp deletion, and 3 bp insertion at different positions in 22 susceptible lines, producing disturbed recognition site for important transcription sites in the intronic region of *Dormancy/auxin associated protein* resulting into loss of signaling in disease resistance pathways. *Dormancy/auxin associated protein* also works as a response protein for signaling molecules important in growth and development of plants. Therefore, disruption in the function of *Dormancy/auxin associated protein* in SMD susceptible genotypes might be resulting in pollen sterility, pollen abortion or seed shriveling/sterility. The Indels and SNP variants are already validated on the resistant and susceptible genotypes of WGRS panel and RIL population (Entire used in this study) respectively. These validated Indel markers are located in the intronic region of the *Dormancy/auxin associated protein* and hinder the recognition sites for RTV1 transcription factor, MYB2 transcription factor, Basic helix-loop-helix (bHLH) transcription factor, and AT-hook transcription factor. These transcription factors regulate the expression of *Dormancy/auxin associated protein* gene. These validated markers could be converted into diagnostic markers and could be used to identify the SMD resistant and susceptible lines for their use in improving SMD resistant pigeonpea cultivars. The identified important genes from this study could also be exploited for gene cloning, functional validation of genes for the viral disease resistance in pigeonpea or other legume species.

## Authors Contribution

**Sagar Krushnaji Rangari:** Writing-original draft; investigation; methodology; formal analysis; validation**. Namita Dube**: formal analysis; software; data curation; writing-review and editing. **Vinay Sharma**: Methodology; formal analysis; writing-review and editing. **Sunil S Gangurde**: Writing original draft; writing-review and editing; formal analysis. **Mamta Sharma**: writing-review and editing; data generation and data curation**. Prakash I Gangashetty**: writing-review and editing. **Rachit K Saxena:** Writing-review and editing, population development. **Abhinav Moghiya**: formal analysis; writing-review and editing. **Vinay K Sharma**: Writing-review and editing, supervision. **K. L. Bhutia**: Writing-review and editing. **Ravi Kant**: Writing-review and editing. **Mahendar Thudi**: Writing-review and editing; formal analysis. **Satheesh Naik SJ:** Writing-review and editing. **Aditya Pratap**: Writing-review and editing. **Girish P Dixit:** Writing-review and editing. **Sean Mayes:** Writing-review and editing. **Manish K Pandey**: Conceptualization; data generation; supervision; methodology; funding acquisition, investigation, writing-original draft, writing-review and editing.

## Supplemental information

**Supplementary table 1: Phenotypic information of parents and bulks**

**Supplementary table 2: sequencing statistics of the genotypes used in marker validation Supplementary table 3: Number of variants chromosome wise**

**Supplementary table 4: Identification of SNPs and Indels in the identified genomic regions (≥99)**

**Supplementary table 5: Identification of Synonymous nonsynonymous SNP Indels**

**Supplementary table 6: Genes identified from the candidate genomic region**

**Supplementary table 7: Resistance genes from identified regions**

**Supplementary table 8: Putative markers selected based on higher p values**

**Supplementary table 9: KASP marker validation on PRILE population Supplementary figures:**

**Supplementary figure 1: Genome-wide SNP index of S-bulk identified for SMD resistance using the ICPL 87119 (Asha) as reference genome**

**Supplementary figure 2: Genome-wide SNP index of R-bulk identified for SMD resistance using the ICPL 87119 (Asha) as reference genome**

**Supplementary figure 3: Genome-wide ΔSNP index identified for SMD resistance using the ICPL 87119 (Asha) as reference genome**

## Data availability statement

All the data generated in the present study is provided in the Supplementary Information and sequencing data deposited as Bioproject ID PRJNA1184965, PRJNA575817, PRJNA383013, PRJNA57990 in NCBI.

