## Supplementary Figure 1 to 3 for "InDels in an intronic region of gene *Ccsmd04* coding for dormancy/auxin-associated protein controls sterility mosaic disease resistance in pigeonpea"

### InDels in an intronic region of gene *Ccsmd04* coding for dormancy/auxin-associated protein controls sterility mosaic disease resistance in pigeonpea

Sagar K Rangari[1,2], Namita Dube[1], Vinay Sharma[1], Sunil S Gangurde[1], Mamta Sharma[1], Prakash I Gangashetty[1], Rachit K Saxena[1,3], Abhinav Moghiya[1], Vinay K Sharma[2], K. L. Bhutia[2], Ravi Kant[4], Mahendar Thudi[5], Satheesh Naik SJ[6], Aditya Pratap[6], Girish P Dixit[6], Sean Mayes[1], Manish K Pandey[1,*]

[1]Center of Excellence in Genomics & Systems Biology (CEGSB), and Center for Prebreeding Research (CPBR), International Crops Research Institute for the Semi-Arid Tropics (ICRISAT), Patancheru, 502324, Hyderabad, India.

[2]Department of Agricultural Biotechnology and Molecular Biology, Dr Rajendra Prasad Central Agricultural University, Pusa, 848125, Bihar, India.

[3]ICAR-Indian Agricultural Research Institute, Dirpai Chapari, 787034, Assam, India.

[4]Department of Genetics and Plant Breeding, Tirhut College of Agriculture (TCA), Dholi, Dr Rajendra Prasad Central Agricultural University, Pusa, 848125, Bihar, India.

[5]College of Agriculture, Family Sciences and Technology, 1005 State University Dr Fort Valley State University, Fort Valley, GA, USA.

[6]ICAR-Indian Institute of Pulses Research (IIPR), Kanpur, Kalyanpur, 208024, Kanpur Uttar Pradesh, India.

ORCID: Manish K Pandey https://orcid.org/0000-0002-4101-6530

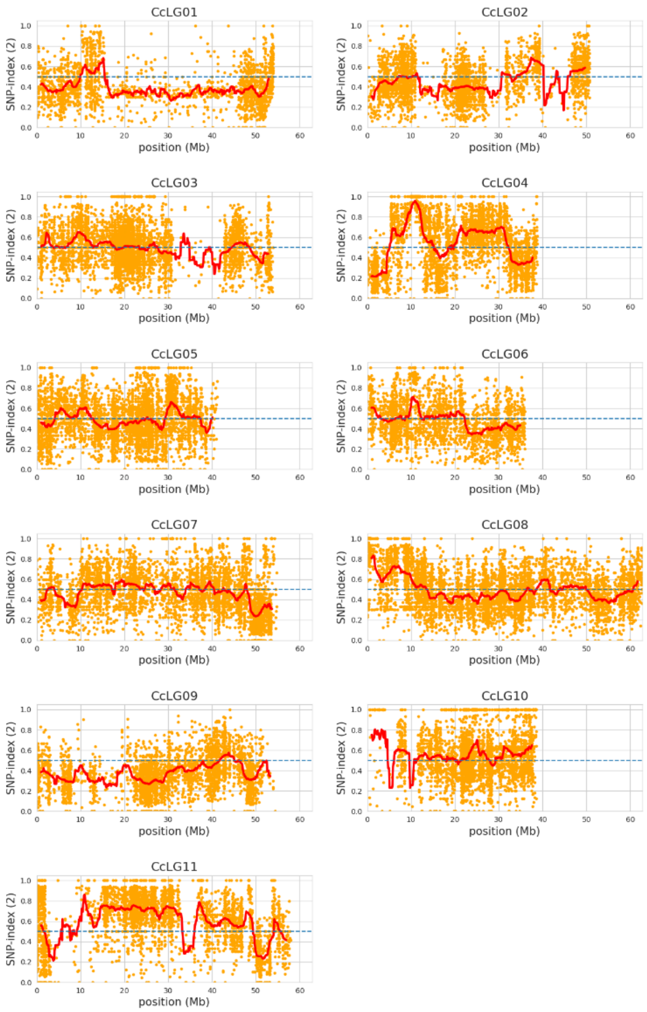


Supplementary Fig 1. Genome-wide SNP index of S-bulk identified for SMD resistance using the ICPL 87119 (Asha) as reference genome.

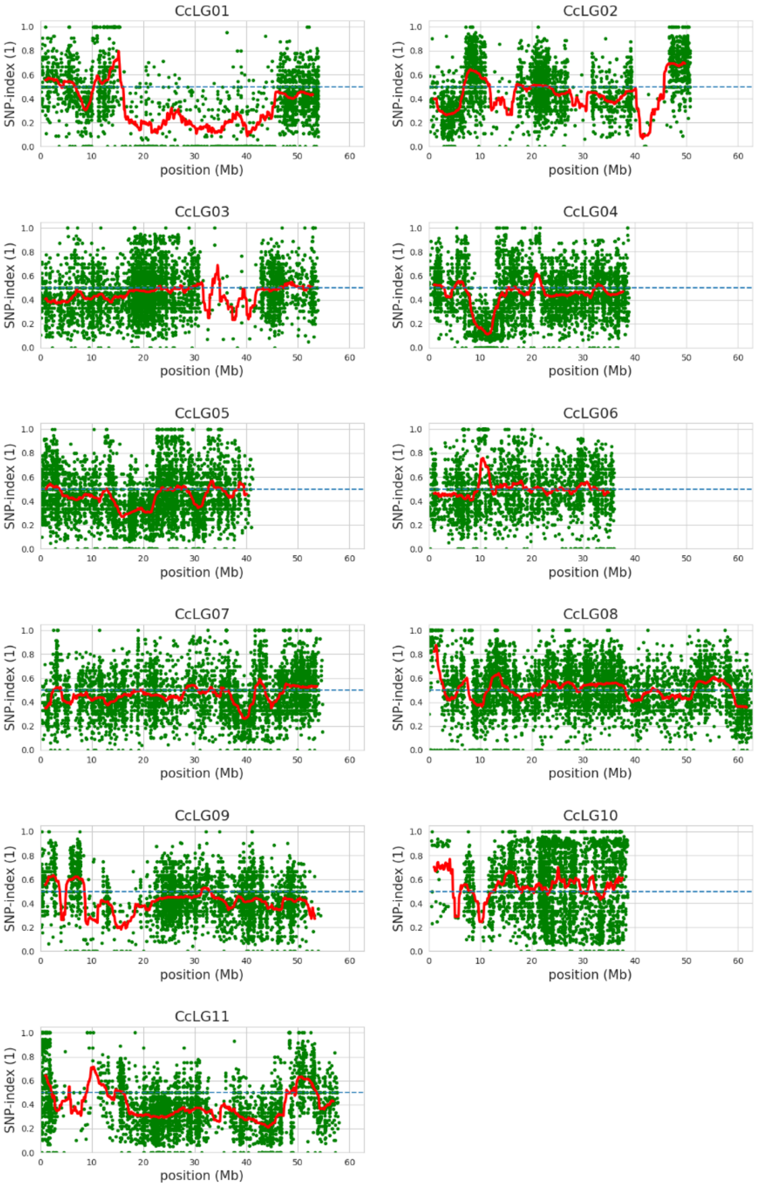
CcLG01
CcLG02
CcLG03
CcLG04
CcLG05
CcLG06
CcLG07
CcLG08
CcLG09
CcLG10
CcLG11
SNP-index (1)
position (Mb)

Supplementary Fig 2. Genome-wide SNP index of R-bulk identified for SMD resistance using the ICPL 87119 (Asha) as reference genome.

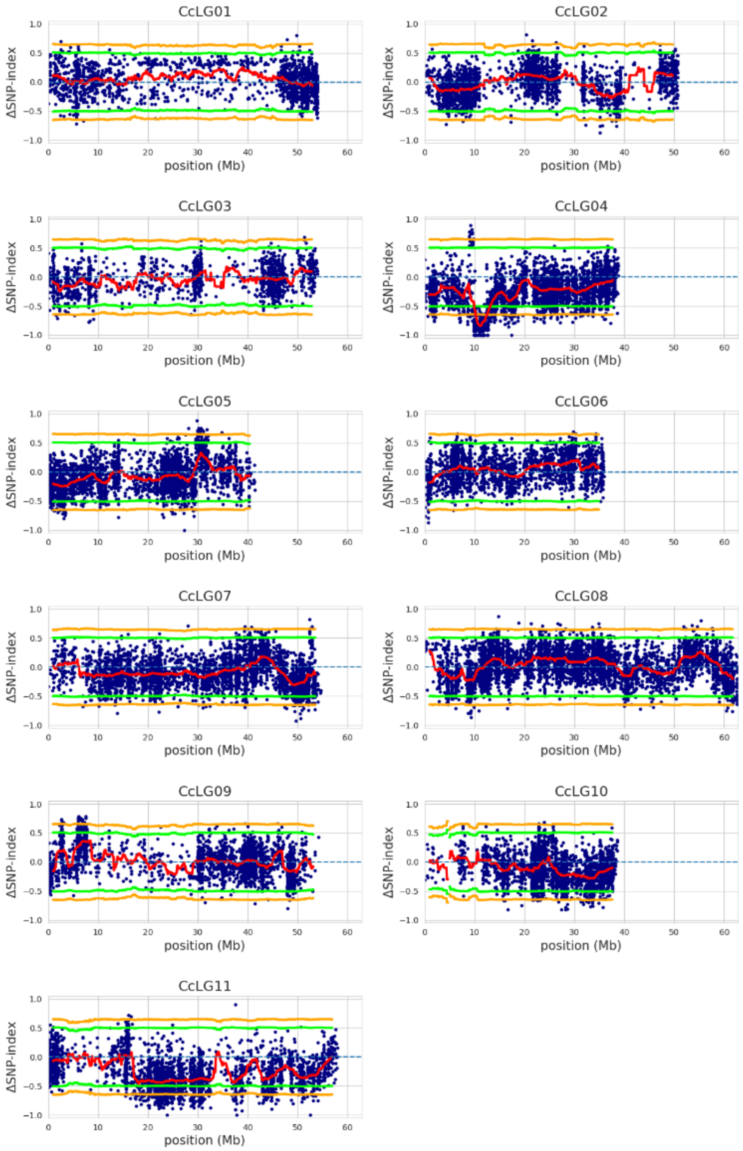
CcLG01
CcLG02
CcLG03
CcLG04
CcLG05
CcLG06
CcLG07
CcLG08
CcLG09
CcLG10
CcLG11
ΔSNP-index
position (Mb)

Supplementary Fig 3. Genome-wide ΔSNP index identified for SMD resistance using the ICPL 87119 (Asha) as reference genome. CI above 99% is indicated by orange line and CI above 95% is indicated by green line
